# Does no-till agriculture limit crop yields?

**DOI:** 10.1101/179358

**Authors:** Matteo Tanadini, Zia Mehrabi

## Abstract

No-till is an agricultural practice widely promoted by governments, development agencies, and agricultural organisations worldwide. However, the costs and benefits to farmers adopting no-till are hotly debated ^1–4^. Using a meta-analysis of unprecedented study size, Pittelkow *et al.*^5^ reported that adopting no-till results in average yield losses of -5.7%, but that these losses can be limited with the added implementation of two additional conservation agriculture practices - crop rotation and crop residue retention, and in dry environments. They claimed that, as a result, resource limited smallholder farmers, that are unable to implement the whole suite of conservation agriculture practices are likely to experience yield losses under no-till. In a re-evaluation of their analysis, we found that they overly biased their results toward showing that no-till negatively impacts yields, and overlooked the practical significance of their findings. Strikingly, we find that all of the variables they used in their analysis (e.g. crop residue management, rotation, site aridity and study duration) are not much better than random for explaining the effect of no-till on crop yields. Our results suggest that their meta-analysis cannot be used as the basis for evidence-based decision-making in the agricultural community.

---

There is never a perfect analysis, and every analysis involves a large number of decisions, some of which are often arbitrary. However, our re-evaluation found four major issues with Pittelkow *et al.*’s meta-analysis, which are fundamental, based on standard statistical theory and best practice. We show how methodologically accounting for these issues produces results that strongly challenge the claims and conclusions of Pittelkow *et al.*

First, we found that Pittelkow *et al*.’s estimates were biased to showing no-till leads to a yield loss. To obtain their -5.7% yield loss figure, they log transformed the ratio of crop yield under no till and conventional till for each experiment, calculated a weighted mean effect of no-till for all these experiments and then backtransformed the result using the exponential function. However, due to Jensen’s Inequality, using the exponential function to backtransform a log-ratio is both biased and inconsistent. In practical terms, this means that the exponentially back-transformed estimates are underestimated (i.e. too small), even when the sample size is very large. The magnitude of the bias depends on the variance of the random variable itself, and in many cases is likely to be negligible. However, for the case of Pittelkow *et al*., there was massive variation in the ratio of yields under no-till and till, which forced large bias in their results towards showing a negative effect of no-till on crop yields. Simply calculating the weighted mean yield ratio in the untransformed scale produces a value of -2.4%, which is around half of the crop yield loss reported by Pittelkow *et al.* (-5.7%).

Second, Pittelkow *et al.* arbitrarily chose to discard observations from their data. They removed all data points further away than 5 standard deviations from the weighted mean, representing about 0.5% of the observations, from their analysis. They did not state any reason for how the threshold of 5 standard deviations was chosen (we are also not aware of any such rule). Because of the long-tailedness of the data, the removal of these observations had an important effect on the results. Including the observations that had been arbitrarily removed from the data leads to a yield ratio of only -1.2%, ∼80% less severe than the figure originally reported. Moreover, removing these observations from the data also did not solve the problem of outliers – it simply created more of them (Supplementary Information A).

Third, we found that the statistical significance testing reported by Pittelkow *et al*. was invalid. To calculate the confidence intervals around, and statistical significance of the effect of no-till on crop yields Pittelkow *et al.* used randomization and bootstrapping techniques. A central assumption of the methods they used is the independence of observations^6^. However, the 5493 experiments in their dataset arose from 609 different studies and so are unlikely to be independent. Our analysis clearly shows that observations gathered in the same study are more similar than observations between studies (Supplementary Information A), which in turn invalidates Pittelkow et al.'s statistical tests. In other words, the statistical significance testing reported in their article should be disregarded. There are established ways to account for this non-independence (Supplementary Information A). However, due to the large sample size of this data set, there is also considerable power to detect statistical significant results, even if practical significance is lacking.

Finally, we found that Pittelkow et al. overlooked the practical significance of their findings. In their paper, Pittelkow *et al.* acknowledged that there was variability in the effect of no-till on yields, and in turn acknowledged that there were statistical differences in the effects of no-till between combinations (or groups) of agricultural practices. However, they did not mention, or visualise, the huge variability in the effects of no-till within each of these groups, nor determine how well their analysis was able to explain this variation. This is problematic because determination of the variation explained by moderators in meta-analysis is a critical component of interpreting the size and statistical significance of their effects^7^. Whilst ignored by Pittelkow *et al.*, an indication of the practical significance of moderators used in a meta-analysis can be assessed graphically, or by using a simple meta-analytic equivalent of the *R*^*2*^ statistic ^7-8^.

In all, forty-three per cent of experiments in Pittelkow *et al.*’s data show the same or higher yields under no-till. Moreover, the majority of yield ratios (e.g. 95% quantiles), lay between a halving in yields (-51%), to a three quarter increase in yields (+74%) under no-till. As shown graphically in Figure 1a, this variation makes it very difficult to make any general claims about negative effects of no-till on crop yield. When we assessed how well the moderators used by Pittelkow et al. (i.e. crop residue management, rotation, site aridity and study duration) helped to explain this large degree of variation, we found that they all performed extremely poorly. As clearly shown in Figure 1b, the cumulative explanatory power of all of the moderators Pittlekow et al. used in their analysis (that is, the combined knowledge crop rotation, residue management, site aridity, and study duration for explaining the effect of no-till on yield) is close to zero, with an *R^2^* of 0.03. Thus, whilst Pittelkow *et al.* stated that crop rotation, residue management and aridity were generally important factors moderating yields under no-till, we found no support for these conclusions.

**Figure 1.**
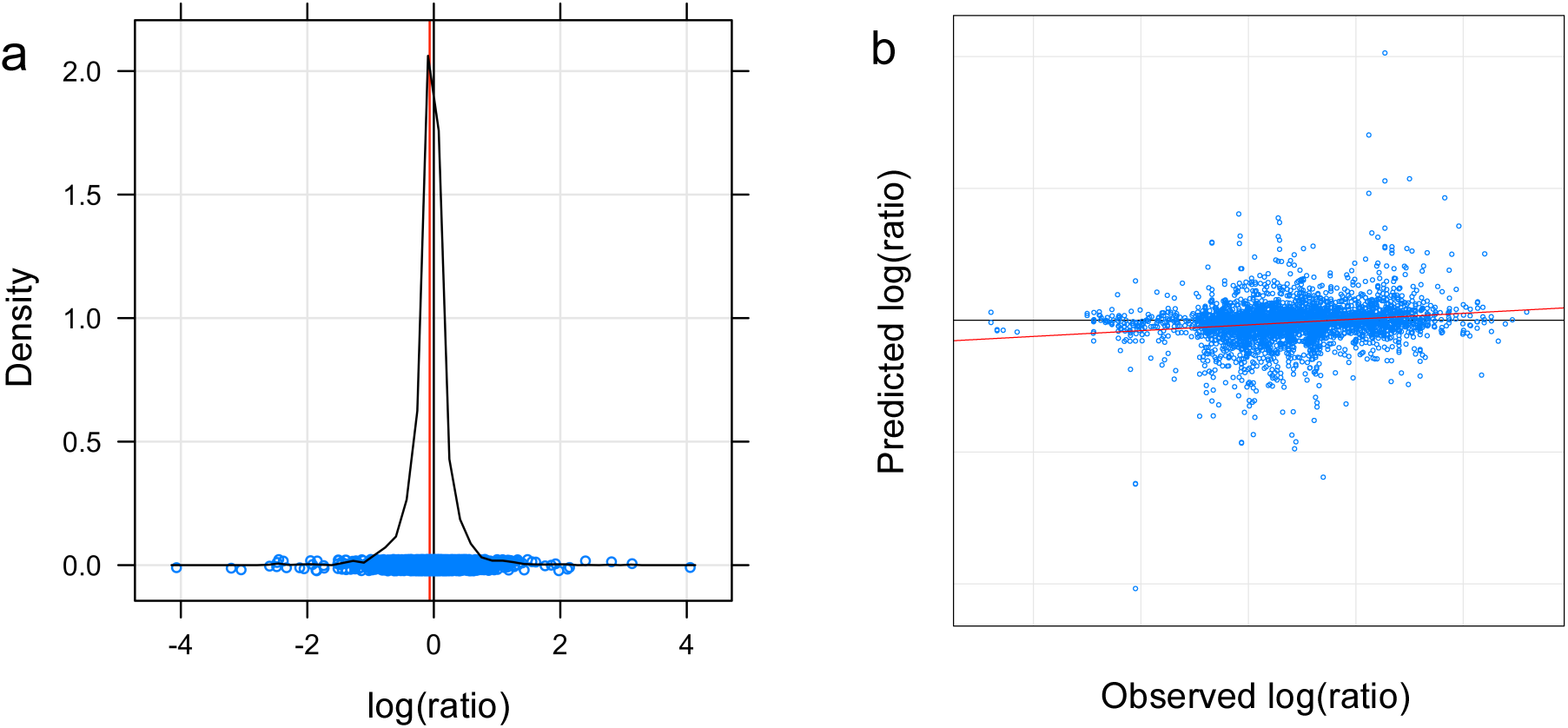
(a) This smoothed histogram (also called a density plot) shows the distribution of the log(ratio) (i.e. the logarithm of the ratio between no-till and till yields). The weighted mean of the data (red line) can hardly be distinguished from the null hypothesis of no difference between yields under no-till and till (black line on zero). (*n* observations= 5492). (b) This graphical *R*^*2*^ scatterplot illustrates the relationship between the predicted and observed values of a linear model including all of the moderators used in the analysis of Pittelkow *et al.* It shows that the crop rotation, crop residue management, aridity and study duration moderators, as used by Pittelkow *et al.* as the basis for their analysis, poorly explain differences in the effect of no-till on yields, with a cumulative explanatory power close to zero (*n* observations = 3753; with a reduction in *n* due to the loss of observations from studies which failed to report on moderators, and so couldn’t be used to assess the interactions between moderators that formed the basis of Pittelkow *et al.*’s main claims in their paper).

The poor association between the moderators used by Pittelkow *et al.* and the effect of no-till on crop yield, suggests it makes little sense to make the general claims that rotation, residue management or site aridity will help increase yields under no-till. It also makes little sense to base further claims on this finding: such that we might expect yield losses to smallholder farmers, such as those in sub-Saharan Africa, due to their inability to implement additional conservation agricultural management practices. However, the applicability of Pittelkow *et al.*’s original analyses to smallholders in sub-Saharan Africa should be questioned anyway, as observations from this region only make up ∼6% of their data set, with the majority of observations (69%) coming from N. America, Europe & New Zealand.

In summary, our analysis suggests that Pittelkow *et al*.’s claim that no-till on average reduces crop yields by 5.7% is incorrect. We found that mean yields under no-till are actually up to 80% less negative than those they reported (-1.2%). This, in addition to the large variation in experimental outcomes, and the poor explanatory power of the variables used by Pittelkow *et al*. in their analysis, strongly challenges the central claims of their paper. Our analysis does not corroborate their claim that farmers will experience yield deficits under no-till, or that rotation and residue management will increase yields under no-till. With the large number of possible factors that could drive differences in yields under no-till, we suggest that much more careful attention be put towards better variable coding, model building, and model interpretation in future if general statements on the outcomes of different management practices under no-till are to be made.

There are of course, other, important reasons why farmers may wish to employ crop residue management and crop rotation alongside no-till in dry environments (e.g. to control pests, soil erosion and soil moisture retention). Similarly, there are other, potential limiting factors to the uptake of no-till agriculture by smallholders in some regions of the world (e.g. access to herbicides, implements)^3-4^. However, we found no support for the claim that yield losses are to be expected for many of the smallholders currently adopting of no-till practices, or that rotation, residue management or dry environments generally increase yields under no till. A wealth of literature has already been published to try and aid biologists in making meta-analyses more methodologically robust^9-11^. In an effort to try and further help future researchers working with large datasets and meta-analysis, we detail methods for dealing with the issue we have outlined here, and many additional others, which did not fit in this response in the Supplementary Information accompanying this article.

## Methods

All methods, advice for dealing with other aspects of reproducibility and open science practice relating to the original Pittelkow *et al.* article, such as data set formatting, code presentation, and transparency of methods; and other guidance on treatment of large datasets, non-normality, non-independence, and practical vs. statistical significance, are described in detail in the Supplementary Information. We note that newer publications^12-13^ following the original Pittelkow *et al.* article do not overcome the problems we have raised here.

## Author contributions

MT & ZM designed the study. MT wrote the first draft of the analysis. ZM wrote the first draft of the paper. Revisions were undertaken by both MT & ZM.

## Footnotes

1 Liu Institute For Global Issues, Institute for Resources, Environment and Sustainability, & Centre for Sustainable Food Systems, University of British Columbia, Vancouver, British Colombia, Canada, V6T 1Z2

